# Age dependency of neurometabolite T_1_ relaxation times

**DOI:** 10.1101/2024.09.30.615917

**Authors:** Saipavitra Murali-Manohar, Helge J. Zöllner, Kathleen E. Hupfeld, Yulu Song, Emily E. Carter, Vivek Yedavalli, Steve C. N. Hui, Dunja Simicic, Aaron T. Gudmundson, Gizeaddis Lamesgin Simegn, Christopher W. Davies-Jenkins, Georg Oeltzschner, Eric C. Porges, Richard A. E. Edden

## Abstract

**Purpose:** To measure T_1_ relaxation times of metabolites at 3T in a healthy aging population and investigate age dependence.

**Methods:** A cohort of 101 healthy adults were recruited with approximately 10 male and 10 female participants in each ‘decade’ band: 18-29, 30-39, 40-49, 50-59, and 60+ years old. Inversion-recovery PRESS data (TE/TR: 30/2000 ms) were acquired at 8 inversion times (TIs) (300, 400, 511, 637, 780, 947, 1148 and 1400 ms) from voxels in white-matter-rich centrum semiovale (CSO) and gray-matter-rich posterior cingulate cortex (PCC). Modeling of TI-series spectra was performed in Osprey 2.5.0. Quantified metabolite amplitudes for total N-acetylaspartate (tNAA_2.0_), total creatine at 3.0 ppm (tCr_3.0_) and 3.9 ppm (tCr_3.9_), total choline (tCho), myo-inositol (mI), and the sum of glutamine and glutamate (Glx) were modeled to calculate T_1_ relaxation times of metabolites.

**Results:** T_1_ relaxation times of tNAA_2.0_ in CSO and tNAA_2.0_, tCr_3.0_, mI and Glx in PCC decreased with age. These correlations remained significant when controlling for cortical atrophy. T_1_ relaxation times were significantly different between PCC and CSO for all metabolites except tCr_3.0_. We also propose linear models for predicting metabolite T_1_s at 3T to be used in future aging studies.

**Conclusion:** Metabolite T_1_ relaxation times change significantly with age, an effect that will be important to consider for accurate quantitative MRS, particularly in studies of aging.

## Introduction

There is substantial clinical interest in understanding cellular and biochemical changes associated with brain aging^1–7^. In vivo magnetic resonance spectroscopy (MRS) offers a unique tool for non-invasively detecting signals from endogenous metabolites in the human brain. MRS is quantitative, in that metabolite levels can be determined based on the amplitude of signals in the in vivo spectrum. While other approaches exist^8,9^, the most common (and consensus) approach to metabolite quantification is to use the endogenous water signal as an internal concentration reference^9,10^. Assuming that the concentration of MR-visible water is a known quantity in different tissue types, it is possible to infer metabolite concentrations from the relative intensity of metabolite signals with respect to the water reference signal. Successive MRS measurements are performed before the signals have fully relaxed to equilibrium in order to maximize signal-to-noise ratio (SNR). Data acquired before the complete recovery of the longitudinal magnetization and acquired at finite echo times (TE) are T_1_- and T_2_-weighted, respectively. Corrections for longitudinal (T_1_) and transverse (T_2_) weighting of both the metabolite and water signals are necessary to yield accurate concentration values. Thus, the extent to which MRS can correctly quantify metabolite levels depends on knowledge of the relaxation times.

Cellular and metabolic changes that occur across the lifespan are of keen interest, both in terms of understanding healthy aging and providing a baseline from which to study disease. Previous MRS studies of the aging human brain have tended to report that levels of NAA and Glu decrease with age^1,2^, while metabolites such as Cho, Cr, and GSH increase^1–4^ with age. One common methodological limitation for MRS studies of aging is that they either do not attempt relaxation correction or apply a single set of relaxation reference values to the whole dataset.

However, such approaches may bias study findings if this age-independent relaxation treatment is not appropriate. There is now substantial evidence that T_2_ relaxation rates change with aging^11–15^. Age-related changes in T_1_ relaxation have not been as extensively studied but would similarly be expected. After inconclusive results from a 1.5T study conducted on two different age groups^16^, there is gradually increasing evidence from another 1.5T study^17^ and our prior 3T study^18^ that T_1_ relaxation times of metabolites change with age. Our recent results demonstrated significant decreases in metabolite T_1_s across the adult lifespan but were limited by a two-point methodology and only considering two metabolites^18^.

In this manuscript, a new prospective cohort (approximately 10 females and 10 males in each decade of adult life from 20s, 30s, 40s, 50s, and 60+) was recruited to measure metabolite T_1_ relaxation times across the lifespan. With the balanced age and sex makeup, this study will fill a current gap by producing age-normed reference values for quantification correction in the future. There is also a growing interest in the relaxation properties of metabolite signals^19^, not just for the purposes of quantification correction, but for information on the specific cellular environments sampled by metabolites^20^.

## Methods

### Participants

A total of 101 participants were recruited for this study at two sites, the Johns Hopkins University School of Medicine (JHU, n = 51) and the University of Florida (UF, n = 50). The age range of the participants was balanced in such a way that it included approximately 10 females and 10 males (5 females and 5 males at each site) from age ranges: 18-29, 30-39, 40-49, 50-59, and 60+ years. All study procedures were approved by a single Institutional Review Board (sIRB) at the Johns Hopkins University School of Medicine, with authority ceded by the University of Florida IRB. Written informed consent was obtained from all participants. A detailed description of participant demographics can be found in Table 1. A previous publication^15^ reports on metabolite T_2_s from the same scan session.

**Table 1:** Participant demographics.

| Variable | 18-29 years | 30-39 years | 40-49 years | 50-59 years | 60+ years |
| --- | --- | --- | --- | --- | --- |
| Sex, # | 10 F, 13 M | 9 F, 10 M | 10 F, 9 M | 9 F, 9 M | 11 F, 9 M |
| Age, Mean (SD),<br>[Min, Max], years | 24.2 (3.3)<br>[18.7, 29.8] | 35.2 (2.8)<br>[31.8, 39.9] | 43.9 (2.9)<br>[40.0, 48.2] | 55.0 (2.9)<br>[50.2, 59.8] | 67.5 (4.1)<br>[60.8, 75.4] |

### Data Acquisition

MRI and MRS scans were performed using a 32-channel head coil on either a 3T Philips dStream Ingenia Elition MRI scanner at JHU or a 3T Philips MR7700 MRI scanner at UF. First, a T_1_-weighted structural MRI (MPRAGE, TR/TE: 8.1/3.7 ms, flip angle: 8°, slice thickness: 1.0 mm, 150 slices, acquisition duration: 2 min 46 sec) was acquired to facilitate voxel placement. Next, an inversion recovery series was acquired using PRESS localization (TE/TR: 30/2000 ms, CHESS water suppression with 115 Hz bandwidth) in two voxels (size 30 (AP) × 26 (RL) × 26 (FH) mm^3^), the white-matter-rich centrum semiovale (CSO) and the gray-matter-rich posterior cingulate cortex (PCC). The series consisted of eight logarithmically-spaced inversion times (TIs): 300; 400; 511; 637; 780; 947; 1148 and 1400 ms. 24 transients were acquired per TI sampled at 2000 Hz with 1024 points. A water reference TI series was acquired with the same parameters, except with only 2 transients per TI and no water suppression applied.

### MRS Data Processing

TI series MRS data were processed using Osprey 2.5.0, an open-source analysis toolbox for MRS data^21^. All processing steps recommended in the MRS community consensus^9^ were followed. Philips raw data were coil-combined, eddy-current-corrected and averaged within each TI on the scanner. The residual water signal was removed using Hankel singular value decomposition (HSVD).

### MRI Segmentation

Osprey-constructed binary masks of the voxels were co-registered to the T_1_-weighted structural images and segmented using SPM12^22^ to yield volume fractions of white matter (fWM), gray matter (fGM), and cerebrospinal fluid (fCSF).

### Spectral Modeling

Spectral modeling was performed for each TI spectrum separately in Osprey 2.5.0 using a custom PRESS TE 30 ms basis set generated using MRSCloud^23^ (https://braingps.mricloud.org/mrs-cloud) for Philips-specific sequence timings and accurate RF pulse shapes. The basis set consisted of 19 metabolite basis functions: ascorbate Asc, aspartate Asp, the methyl group of creatine CrCH_3_ at 3.0 ppm, methylene group of creatine CrCH_2_ at 3.9 ppm, gamma-aminobutyric acid GABA, glycerophosphocholine GPC, glutathione GSH, glutamine Gln, glutamate Glu, lactate Lac, myo-inositol mI, acetyl moiety of NAA NAA_ace_ at 2.01 ppm, aspartyl moiety of NAA NAA_asp_, acetyl moiety of N-acetyl aspartyl glutamate NAAG_ace_ at 2.04 ppm, aspartyl and glutamate moiety of NAAG NAAG_asp_glu_, phosphocholine PCh, phosphorylethanolamine PE, scyllo-inositol sI, taurine Tau; 5 macromolecule basis functions: MM09, MM12, MM14, MM17, MM20; and 3 lipid basis functions: Lip09, Lip13, Lip20. Macromolecules and lipids were included in modeling as parameterized Gaussian basis functions.

In line with other linear combination modeling software, Osprey only allows positive coefficients for basis functions in the spectral model. This presents a challenge for modeling TI series spectra in which signals have a polarity that depends on the degree of longitudinal relaxation of the z-component during the inversion delay TI. For example, in short-TI spectra, metabolite signals appear with negative polarity whereas macromolecule signals have already recovered past the null-point and appear with positive polarity. It is convenient for modeling to phase these spectra so that the metabolite signals are positive – therefore macromolecule and lipid basis functions were inverted. In TI-511 spectra, macromolecule signals again appear with positive polarity, and metabolite signals are still inverted but close to the null point. These spectra were modeled with their natural polarity, so metabolite basis functions were inverted. In TI-637 spectra, the metabolite signals are very close to the null point with all but the strongest signals well below the noise level. These spectra were modeled with a reduced metabolite basis set (including only positive tCr_3.9_, and negative tNAA_2.0_, tNAA_asp_, mI, Glx, and tCr_3.0_ functions) to avoid over-fitting. At long TIs, both metabolite and macromolecular signals have positive phase and modeling is straightforward. Overall, this approach can be summarized as modeling spectra phased so that the dominant component (i.e. metabolites in long- and short-TI spectra and macromolecules in mid-TI spectra) is positive, and adjusting the polarity of basis functions of other signals as appropriate. This provided better initialization for modeling variables which results in better optimization outcomes. Most TIs were modeled with a spline baseline knot spacing of 0.4, the default for ‘typical’ short-TE spectra at 3T. Mid-TI spectra (TI 511 and 637 ms) were modeled with a more flexible baseline with a knot spacing of 0.2, and without lineshape convolution beyond Gaussian and Lorentzian broadening. Following spectral modeling of the metabolite TI-series, basis function amplitude factors were extracted for T_1_ modeling. Metabolite signal amplitudes for TIs 300, 400, 511 and 637 ms were multiplied by –1 as appropriate to correct for modeling of inverted spectra and/or modeling with inverted basis functions. TI-series water reference spectra were processed to have positive phase and modeled using Osprey with a simulated water basis function. Water amplitudes for T_1_ modeling were extracted and polarities were corrected based on observing the phase inversion of the first point of the raw FIDs.

### T_1_ Relaxation Modeling

For modeling T_1_ relaxation times of the metabolite signals, 6 signals were considered: tNAA_2.0_; tCr_3.0_; tCr_3.9_; mI; and Glx (Glu + Gln). Linear combination model metabolite amplitudes from the TI spectra for every subject were fitted in MATLAB R2022a using lsqcurvefit to the inversion recovery equation:

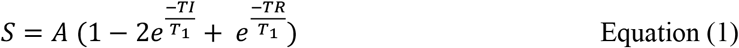

with modeled parameters T_1_ and *A*, an amplitude factor. In order to calculate the T_1_ relaxation time of tissue water, as distinct from CSF water, the water signal amplitudes were modeled using a two-pool inversion recovery model for the water signal *S*_water_:

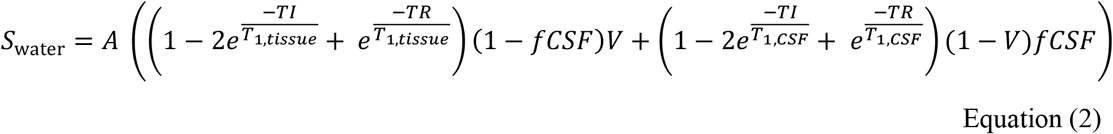

where *A* is an amplitude factor scaling constant, *T*_1,*tissue*_ is the tissue water T_1_ relaxation time, *T*_1,*CSF*_ is the CSF water T_1_ relaxation time, and *V* is a correction factor to account for the differential concentration of MR-visible water between tissue and CSF equal to 0.4211 (i.e. 40/(40+55), with 55 mM being the water concentration in CSF, and 40 mM being an average water concentration estimate in gray and white matter). *A* and *T*_1,*tissue*_ were unconstrained whereas *T*_1,*CSF*_ was constrained between 1000 and 5000 ms.

### Statistical Analyses

All statistical analyses were performed in R 4.3.2^24^ within RStudio^25^ [and followed the identical approach used in our prior report on metabolite T_2_s from the same participants^15^]. The correlation between T_1_ relaxation time and age for each metabolite signal was calculated for each voxel separately. Since multiple variables did not satisfy the Pearson correlation normality assumption (Shapiro test *p* < 0.05), non-parametric Spearman correlations are reported. In order to account for multiple comparisons, Benjamini-Hochberg false discovery rate (FDR) correction was applied to the p-values. A linear model was calculated for each signal with T_1_ as the outcome variable and age-minus-30 as the predictor: *T*_1_ = *β*_*o*_ + *β*_1_ × (Age − 30). The intercept from this model represents the predicted T_1_ at 30 years of age, and the slope represents the change in T_1_ for each year of age. For significant correlations between T_1_ and age, additional analyses were performed controlling the linear models for cortical atrophy: *T*_1_ = *β*_*o*_ + *β*_1_ × (Age − 30) + *β*_2_ × Tissue, using the gray matter tissue fraction (i.e. Tissue = *f*_*GM*_/(*f*_*GM*_ + *f*_*WM*_) to represent the expected loss of cortical GM with age. Two-sample t-tests were used to assess possible sex differences in T_1_ relaxation times in each voxel. Finally, in order to test for differences in T_1_ relaxation times between the voxels, paired t-tests were performed, followed by correction for multiple comparisons.

## Results

### Data Quality

CSO and PCC voxel locations are shown in Figure 1. Voxel tissue fractions were: for CSO, fGM 0.24 ± 0.07, fWM 0.70 ± 0.10, and fCSF 0.06 ± 0.04; and for PCC, fGM 0.58 ± 0.06, fWM 0.25 ± 0.06, and fCSF 0.17 ± 0.07. Four datasets were excluded: one due to motion (49-year-old male), one due to an acquisition error with wrong inversion offset (30-year-old female) and one due to poor linewidth (66-year-old male). For one 19-year-old male, the PCC voxel was placed in the wrong location and that dataset was also excluded. Representative median inversion recovery series spectra from CSO and PCC are shown in Figure 2. T_1_ relaxation curves for the 6 metabolite signals and tissue water are shown in Figure 3 for both CSO and PCC voxels from the same subject. Mean R^2^ values for all the metabolites and water across all the subjects were ≥ 0.95 as a measure of goodness of fit for the T_1_ exponential curves, except tCr_3.9_ which had mean R^2^ = 0.89. Table 2 provides mean values and standard deviations (grouped by age) of T_1_ relaxation times for the 7 signals from CSO and PCC.

**Table 2:** Mean T_1_ relaxation times in milliseconds for metabolites and tissue water grouped by age. Standard deviations are provided in parentheses.

| CSO Voxel |  |  |  |  |  |
| --- | --- | --- | --- | --- | --- |
| Metabolite | 18-29 years | 30-39 years | 40-49 years | 50-59 years | 60+ years |
| tNAA<sub>2.0</sub> | 1112<br>(45) | 1083<br>(45) | 1089<br>(40) | 1070<br>(52) | 1036<br>(44) |
| tCr<sub>3.0</sub> | 1045<br>(54) | 1037<br>(67) | 1040<br>(55) | 1032<br>(39) | 1006<br>(53) |
| tCr<sub>3.9</sub> | 780<br>(35) | 770<br>(44) | 779<br>(34) | 768<br>(26) | 773<br>(35) |
| tCho | 949<br>(38) | 941<br>(44) | 945<br>(36) | 948<br>(29) | 934<br>(43) |
| mI | 949<br>(26) | 969<br>(54) | 949<br>(34) | 961<br>(25) | 943<br>(41) |
| Glx | 1352<br>(201) | 1369<br>(220) | 1287<br>(113) | 1307<br>(172) | 1389<br>(361) |
| Tissue Water | 783<br>(91) | 742<br>(71) | 747<br>(45) | 778<br>(55) | 808<br>(110) |
| PCC Voxel |  |  |  |  |  |
| Metabolite | 18-29 years | 30-39 years | 40-49 years | 50-59 years | 60+ years |
| tNAA<sub>2.0</sub> | 1091 | 1054 | 1051 | 1014 | 998 |
|  | (50) | (41) | (45) | (38) | (46) |
| tCr<sub>3,0</sub> | 1054 | 1054 | 1034 | 1034 | 1022 |
|  | (50) | (50) | (49) | (26) | (47) |
| tCr<sub>3,9</sub> | 737 | 739 | 753 | 748 | 751 |
|  | (32) | (38) | (41) | (22) | (45) |
| tCho | 965 | 964 | 970 | 968 | 963 |
|  | (53) | (54) | (50) | (29) | (45) |
| mI | 1016 | 1018 | 994 | 984 | 994 |
|  | (46) | (59) | (25) | (23) | (42) |
| Glx | 1157 | 1179 | 1091 | 1072 | 1078 |
|  | (78) | (101) | (130) | (81) | (66) |
| Tissue Water | 965 | 971 | 965 | 1008 | 991 |
|  | (113) | (155) | (116) | (172) | (92) |

**Figure 1:**
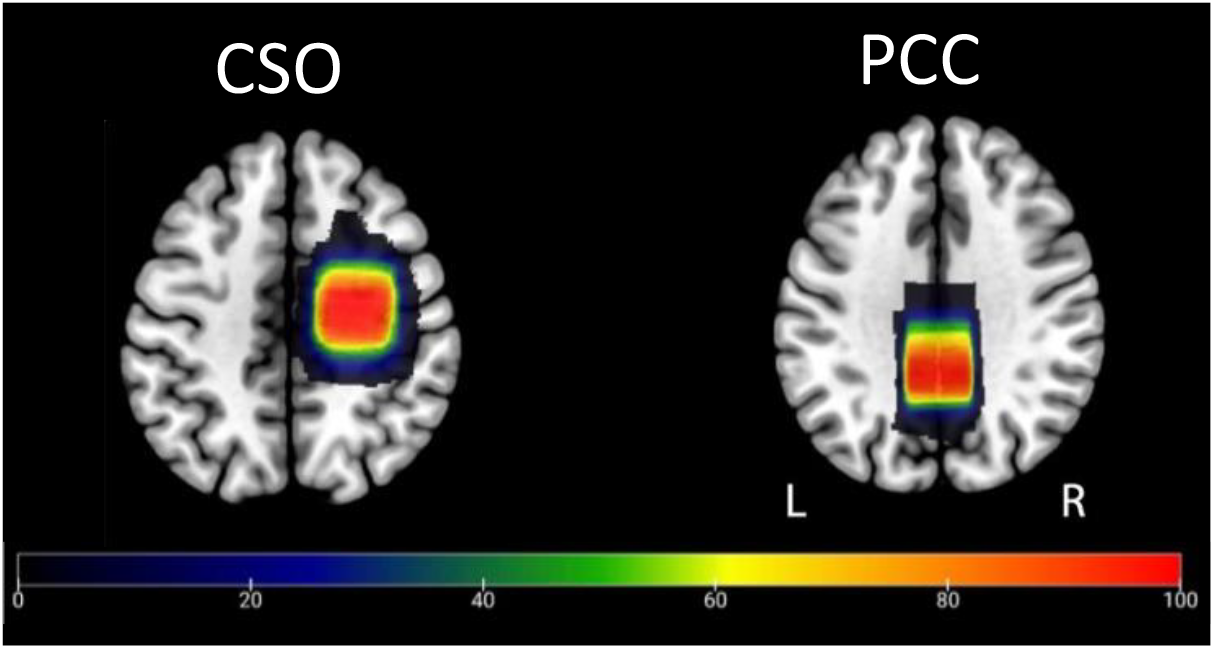
Voxel placement for TI series data acquisition. Voxels of size 30 × 26 × 26 mm^3^ were placed in the CSO and PCC regions. This figure illustrates the areas of overlap in the voxel placements across subjects. The native space binary voxel mask from each participant was normalized to standard MNI space and overlaid onto the spm152 template. The color bar indicates the number of subjects overlapped, from 0 (blue) to all subjects (red).

**Figure 2:**
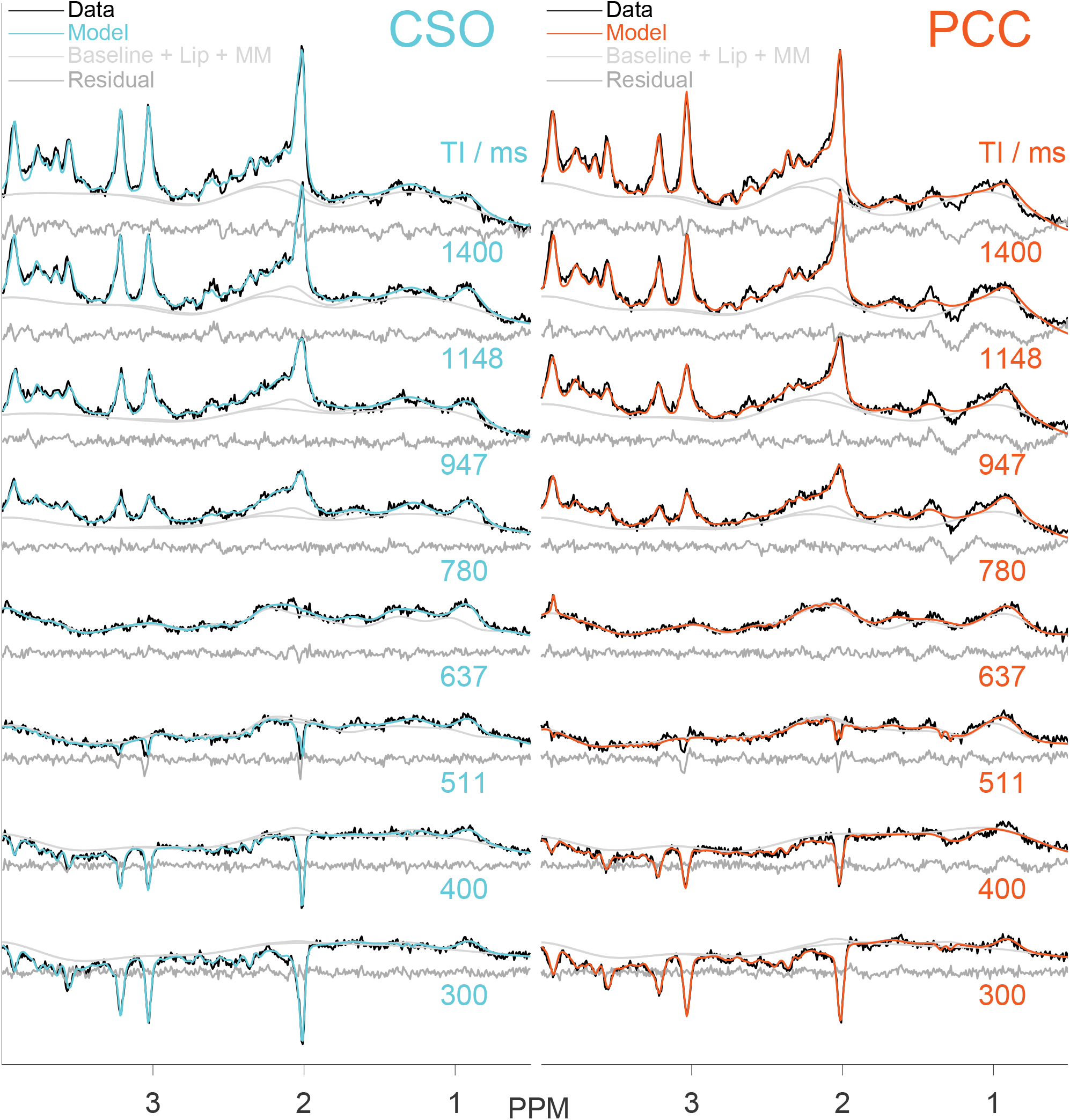
Inversion recovery PRESS spectra, models and fit residuals from a representative subject (47-year-old-female) for all 8 TIs from the CSO and the PCC regions. This subject was determined from the median value of average R^2^ of T_1_ curve model fit for all 6 metabolites and tissue water.

**Figure 3:**
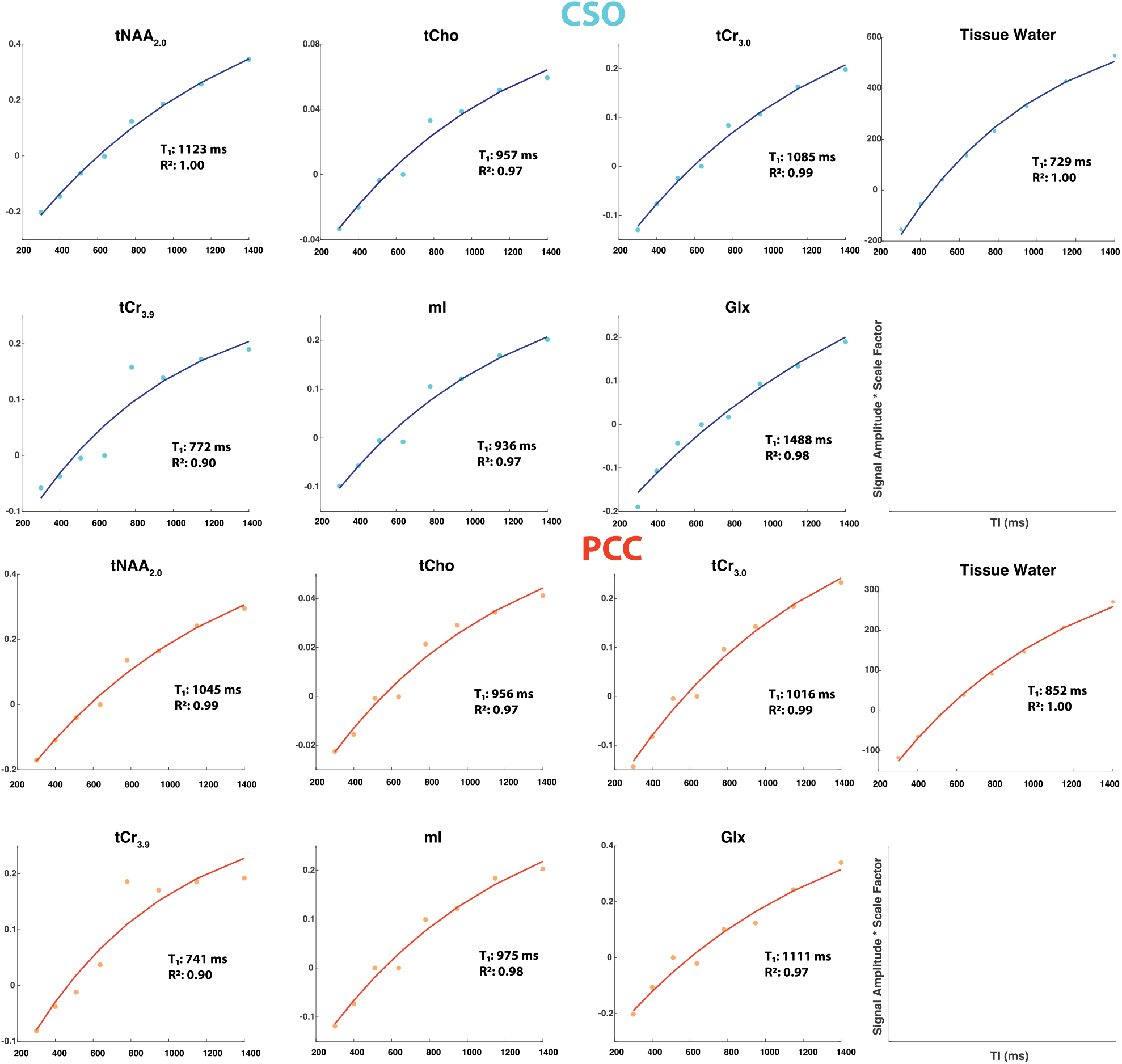
T_1_ relaxation curves for 6 metabolite signals and tissue water from the same subject represented in Figure 2. Points represent the metabolite amplitudes modeled at each TI and solid lines represent the T_1_ relaxation model of best fit as calculated using the inversion recovery signal equation given in Equation 1. T_1_ relaxation times for each metabolite for this subject are provided in the plot along with the R^2^ values as a measure of goodness of fit for the curves.

### T_1_ Correlations with Age

Correlation plots between T_1_ relaxation times and age are presented in Figure 4. Spearman correlation coefficients and FDR-corrected *p*-values are given in Table 3. T_1_ relaxation times decreased significantly with age for tNAA_2.0_, tCr_3.0_, mI and Glx in PCC, whereas only for tNAA_2.0_ in CSO. Among the significant correlations (*p* < 0.05, FDR corrected), Spearman coefficients ranged from –0.28 for mI in PCC to –0.61 for NAA in PCC. Tissue water T_1_ did not correlate significantly with age in CSO or PCC. When controlling for cortical atrophy, correlations of T_1_ with age remained significant for each of the four metabolite signals in PCC and for tNAA_2.0_ in CSO. No sex differences in T_1_ relaxation times of metabolites or tissue water were observed for either region.

**Table 3:**
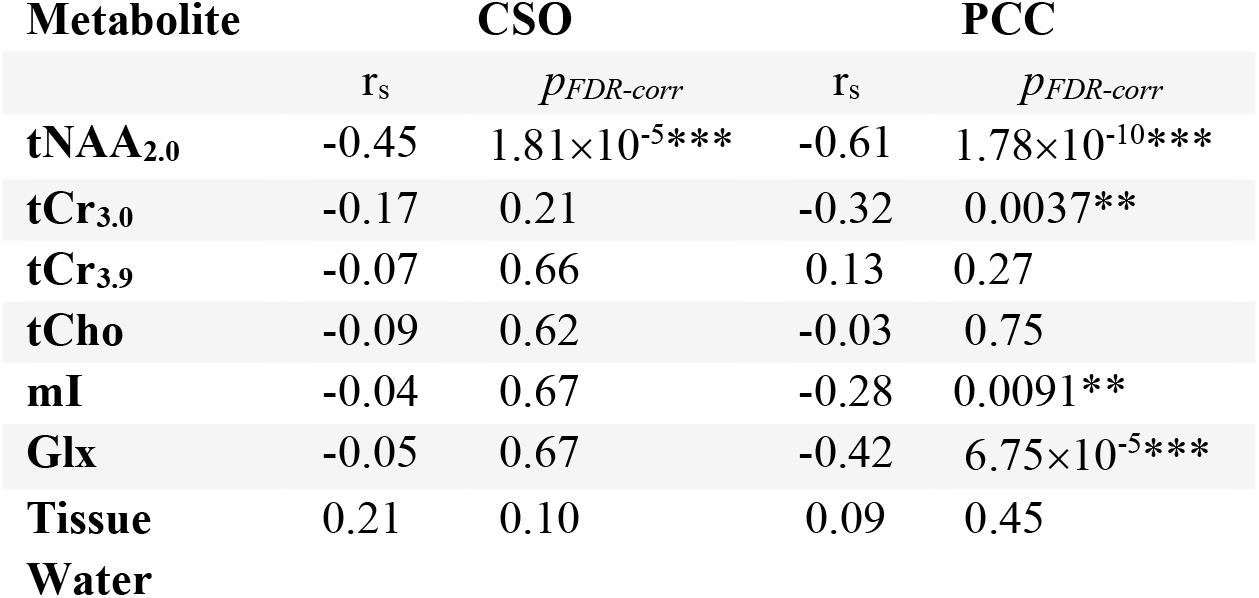
Spearman correlation coefficients (r_s_) of T_1_ with age and FDR-corrected *p*-values. (* represents *p* < 0.05, ** represents *p* < 0.01, *** represents *p* < 0.001) for CSO and PCC.

**Figure 4:**
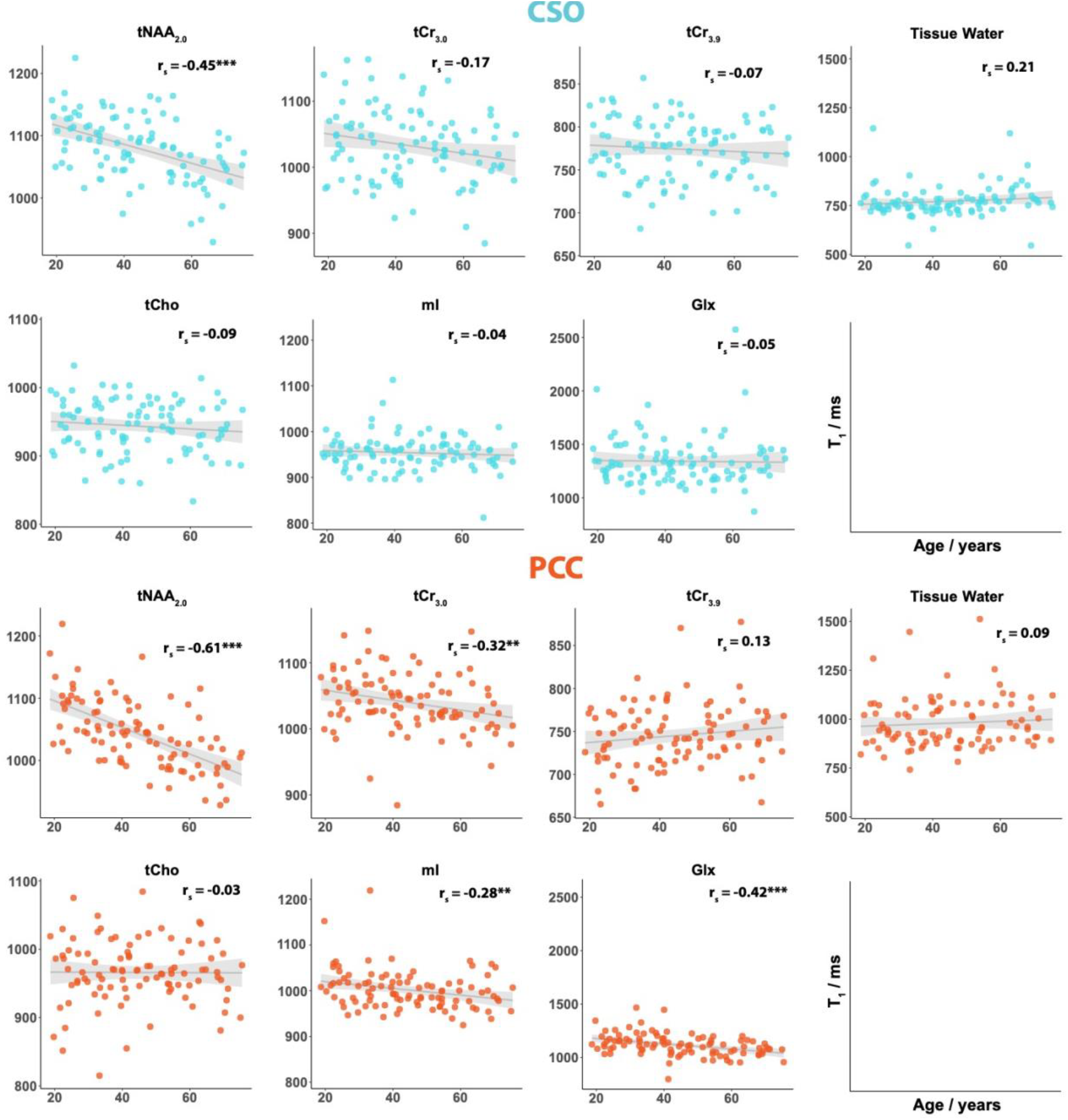
Age-T_1_ correlation plots are shown for all 6 metabolites and tissue water from both CSO (blue) and PCC (orange). Spearman correlation coefficient for each plot is indicated in the top-right corner with FDR-corrected *p*-value for the correlation \**p* < 0.05, \*\**p* < 0.01, \*\*\**p* < 0.001. The gray line and shading represent the linear model and 95% confidence interval for the model using age to predict T_2_ (produced using the *geom_smooth* function in R).

T_1_ can be predicted for each metabolite from the linear model *T*_1_ = *β*_*o*_ + *β*_1_ × (Age − 30). Table 4 gives the T_1_-intercept at 30 years of age (*β*_*o*_), and the slope (*β*_1_) which represents the change in predicted T_1_ for each year of life. Slopes are only reported for the cases where T_1_ was significantly correlated with age. These linear models are represented as the gray trendlines in Figure 4.

**Table 4:** Linear model coefficients, namely intercept and slope, for estimating T_1_ relaxation times for a given age using T_1_ = β_0_ + β_1_*(Age-30). Slope is provided only for those metabolites where T_1_ was significantly correlated with age; slope represents change in predicted T_1_ with each year of life beyond 30 years of age. Intercept values correspond to predicted T_1_ values for Age = 30 years.

| Metabolite | CSO |  | PCC |  |
| --- | --- | --- | --- | --- |
| | Intercept | Slope ($\beta_1$) | Intercept | Slope ($\beta_1$) |
| tNAA<sub>2,0</sub> | 1101 | -1.52 | 1074 | -2.14 |
| tCr<sub>3,0</sub> | 1043 |  | 1051 | -0.75 |
| tCr<sub>3,9</sub> | 777 |  | 740 |  |
| tCho | 947 |  | 966 |  |
| mI | 956 |  | 1012 | -0.74 |
| Glx | 1346 |  | 1152 | -2.46 |
| Tissue Water | 763 |  | 970 |  |

### T_1_ Differences by Voxel

Box plots comparing T_1_ relaxation times for the CSO and PCC voxels are shown in Figure 5. They were significantly different between the voxels for all metabolites, except tCr_3.0_ and tissue water. T_1_ values were higher for tNAA_2.0_, tCr_3.9_ and Glx in CSO, but higher for tCho, mI and tissue water in PCC.

**Figure 5:**
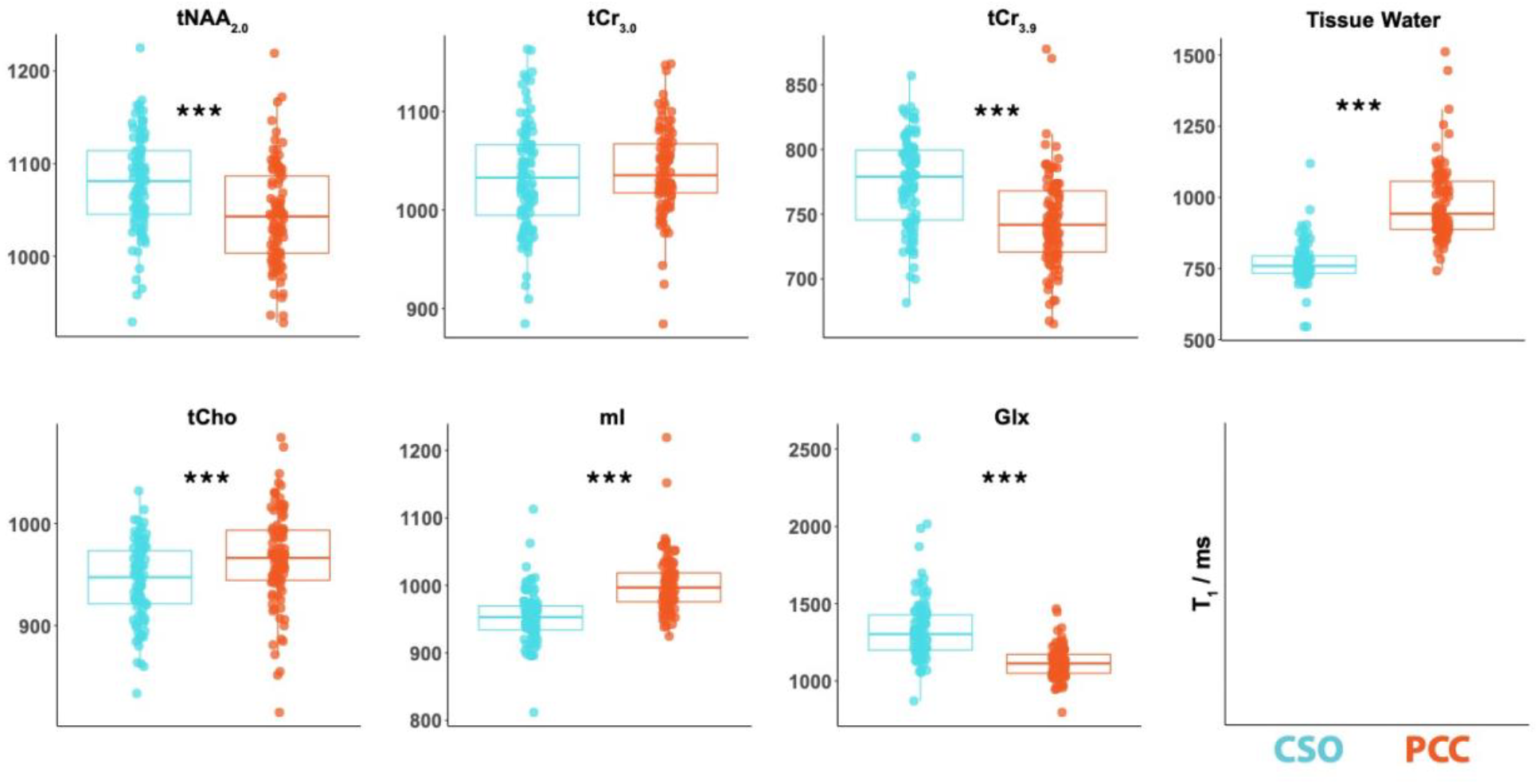
Differences in T_1_ relaxation times between CSO (blue) and PCC (orange). Each datapoint for CSO and PCC represents one participant. The asterisks indicate the statistical significance of the FDR-corrected *p*-value for the paired t-test, \**p* < 0.05, \*\**p* < 0.01, \*\*\**p* < 0.001.

## Discussion

This study investigated changes in T_1_ relaxation times of metabolites and tissue water across the adult lifespan, with participants between 18 and 75 years of age. This cohort of 101 subjects is by far the largest dataset to analyze metabolite T_1_ changes with age. Data were acquired in two regions, WM-rich CSO and GM-rich PCC, which enabled an examination of the differences between WM and GM tissues. The T_1_ relaxation times of most metabolites decreased significantly with age in PCC. However, in CSO, only tNAA_2.0_ decreased significantly with age. T_1_ relaxation times of tissue water did not change significantly in CSO or PCC, though showed a strong positive trend with age in CSO and a weak positive trend in PCC.

To date, few studies^16–18^ have investigated the impact of aging on metabolite T_1_ relaxation. Early research performed at 1.5T was inconsistent with one study showing no changes^16^ and one showing decreases with age^17^ for tNAA_2.0_ and tCr_3.0_ in CSO. In the current 3T study, T_1_ relaxation times decreased significantly with age in PCC for the singlets, tNAA_2.0_ and tCr_3.0_, as well as the multiplets, mI and Glx. In the CSO, significant decreases were only observed for tNAA_2.0_. These results, using a more sophisticated measurement protocol, reproduce and build upon our previous findings^18^ of decreasing metabolite T_1_ with age for tNAA_2.0_ and tCr_3.0_.

There is a more substantial quantitative imaging literature on trends in water T_1_ relaxation times with age. One such study showed significant increases in T_1_ relaxation times of tissue water across various subcortical, cortical and cerebellar brain regions^26^ in a cohort aged between 18 and 78 years old. Another study of 211 subjects aged between 20 and 89 years showed positive correlations between age and water T_1_ in the frontal WM, globus pallidus and genu of the corpus callosum, whereas showed negative linear correlations subcortically in the thalamus, head of the caudate nucleus and putamen^27^. Water T_1_s in the brain are generally thought to be influenced by myelination, iron deposition, and water content. These three factors have been shown to vary with age – myelin and water content decrease with age^27,28^, whereas iron accumulates with age^26,27^. Demyelination prolongs T_1_ relaxation times^29,30^, whereas reduction in water content and increase in iron content reduces T_1_ relaxation times^31,32^. In the current study, the T_1_ of tissue water increases (but not to a significant degree), and more strongly in CSO, possibly reflecting the demyelination associated with aging^33,34^. The difference between PCC and CSO could also reflect the greater influence of iron in GM regions^27^. Metabolites are predominantly intracellular, absent from CSF and present at different levels in different cell types. While the localization of metabolites and water differs, the underlying mechanisms of T_1_ relaxation will be similar but may be weighted differently. The general trend of metabolite T_1_s to decrease with age (whereas water T_1_s tends in general to increase) suggests that the reduction in water content intracellularly and the increase in iron content may have a relatively greater impact on metabolite signals than water.

Prior literature comparing T_1_ relaxation times in GM and WM reveals distinct patterns depending on the metabolite, method, and magnetic field strength used in the measurements. NAA generally exhibits longer T_1_ in WM compared to GM, as observed in the current study, with similar trends reported at 7T^35,36^ and at 1.5 T^37^. However, some studies at 3T^38^ and at 1.5T^39^ reported slightly higher NAA T_1_s in GM than in WM. No significant difference was found in tCr_3.0_ values between WM and GM; results were in line with most previous findings at 1.5T^37^, 7T^35,36^, 4.1T^40^ and 9.4T^41^. However, both higher tCr T_1_ relaxation times at 1.5T^39^, 3T^38^ and lower times at 1.5T^42^ have been reported. Where tissue differences in tCho T_1_s are reported, they tend to be higher in GM at 3T^37,38^ and at 7T^36^, in agreement with the current study. However, they are not always observed^35,39–41^ and the opposite result has been reported at 1.5T^42^. Glx has higher T_1_ values in WM compared to GM in the current study. Results agree with a 7T study ^36^ but disagree with a 3T one ^38^. Findings on T_1_ of mI are inconsistent with higher T_1_s in GM seen in this study, at 3T^37^ and at 9.4T^41^ and lower GM mI T_1_ values reported in a study at 1.5T^42^. Tissue water T_1_ is generally shorter in WM, as indicated in the current study (CSO vs PCC), and in agreement at 3T^37^ and 7T^36^ as well as several non-MRS methods^43–45^. As discussed above, this is likely driven by the higher myelin content in the CSO ^46–48^, accelerating relaxation processes and resulting in shorter T_1_ times^49,50^. It is particularly interesting that one set of metabolites (tNAA, tCr, and Glx) show higher T_1_s in CSO and another (tCho and mI) lower. In terms of their roles and cell type, tNAA and Glx are more closely associated with neuronal energy and neurotransmission metabolism whereas tCho and mI are more associated with the glial compartment^51^. Overall, the variability in metabolite concentrations between WM-rich CSO and GM-rich PCC across these studies reflects the inherent differences in brain tissue composition, the influence of magnetic field strength, and measurement techniques.

This study has a number of limitations. Firstly, modeling of the TI series data was challenging, especially in TIs that are close to metabolite null-points since these spectra had larger MM signals than metabolite signals. The lack of prior knowledge in MM makes it difficult to clean the spectra. Given that our focus here was metabolite T_1_s, we relied upon a spline baseline to model much of the MM contribution to the spectrum, especially at TI 511 and 637 ms, where a more flexible spline baseline was used. Methodologically, it is not ideal to use different model settings for different spectra within a dataset but given the drastically different character of these mid-TI spectra, it was necessary to give plausible spectral models and metabolite amplitudes that were coherent with the inversion-recovery curve established by the long- and short-TI extremes. Reliance upon spline and parametrized MM signals has the potential for overfitting, compared to e.g. using acquired MM background spectra, but no multi-TI reference dataset for MM spectra is available^41^. Secondly, stemming from the limited resolution of MRS, is whether the changes in CSO and PCC reflect general changes in WM and GM tissue respectively, or whether they arise from tissue sampling drift with age. We have attempted to address this limitation using a statistical covariate, but this relies upon the segmentation of T_1_-weighted structural MRI which might itself be biased as water T_1_s change with age and image contrast diminishes. Thirdly, the coverage of this study is limited to just two regions (which may or may not be more broadly representative of cortical GM and WM regions); future work should extend this study to other brain regions. One final limitation is the difficulty of modeling the inversion series spectra reliably, which is reflected in the varied conclusions in prior literature^16–18^ discussed above. It is likely that 2D modelling^52–55^ of the spectra is a more parsimonious and SNR-efficient approach for such datasets. Knowledge can then be shared across TI points, for example, the lineshapes strongly established in long-TI spectra will be constrained for low-SNR mid-TI spectra. However, implementation of a 2D modeling framework is itself not trivial and raises a number of additional decisions about how and whether to share parameters across TIs.

## Conclusion

We report decreasing metabolite T_1_ with age from a large-cohort MRS measurement. This was seen more consistently in the WM-rich CSO than the GM-rich PCC. These results reflect under-studied changes in the cellular microenvironment in the human brain with healthy aging attributed to factors such as myelination, iron deposition and reduced water content. We also provide linear models to predict age-dependent metabolite T_1_s for quantification relaxation correction in future MRS studies.

## Abbreviations

^1^H: Proton
Cho: Choline
Cr: Creatine
CSO: Centrum semiovale
GM: Gray matter
MRS: Magnetic Resonance Spectroscopy
MM: Macromolecule
NAA: N-acetyl aspartate
PCC: Posterior cingulate cortex
PRESS: Point resolved spectroscopy
SNR: Signal-to-noise ratio
TI: Inversion time
WM: White matter.

## Acknowledgement

This work was supported by NIH grants R01 EB016089, R01 EB023963, R00 AG062230, R21 EB033516, K99 AG080084, K00AG068440, and P41 EB031771.

## Author Contributions

SMM processed and modeled all data, conducted all statistical analyses, wrote the manuscript and prepared figures. HJZ contributed to MRS data processing and to the Osprey code for the analysis. KEH contributed to protocol development and contributed to the data analysis and statistical analysis pipeline. CDJ contributed to the Osprey modeling of the data. YS and EC made contributions to data collection. VY reviewed all structural scans to assess data quality and check for incidental findings. ATG and GLS contributed to the manuscript writing. DS contributed a figure. GO, EP, and RAEE designed the project and led the interpretation of the results. SCNH set up the scan protocol and oversaw data quality control. All authors contributed to manuscript revision and approved the final version of the manuscript.

